# Scaling of organ masses in mammals and birds: phylogenetic signal and implications for metabolic rate scaling

**DOI:** 10.1101/694067

**Authors:** Andrzej Antoł, Jan Kozłowski

## Abstract

The persistent enigma of why the whole-body metabolic rate increases hypoallometrically with body mass should be solved on both the ultimate and proximate levels. The proximate mechanism may involve hyperallometric scaling of metabolically relatively inert tissue/organ masses, hypoallometric scaling of metabolically expensive organ masses, a decrease in mass-specific metabolic rates of organs or, more likely, a combination of these three factors. Although there are in data in the literature on the scaling of tissue/organ masses, they do not take phylogenetic information into account. Here, we analyse the scaling of tissue/organ masses in a sample of 100 mammalian species and 22 bird species with a phylogenetically informed method (PGLS) to address two questions: the role of phylogenetic signal in organ/tissue size scaling and the potential role of organ/tissue mass scaling in interspecific metabolic rate scaling. Strong phylogenetic signal was found for the brain, kidney, spleen and stomach mass in mammals but only for the brain and leg muscle in birds. Metabolically relatively inert adipose tissue scales isometrically in both groups. The masses of energetically expensive visceral organs scale hypoallometrically in mammals, with the exception of lungs, with the lowest exponent occurring for the brain. In contrast, only brain mass scales hypoallometrically in birds, whereas other tissues and organs scale isometrically or almost isometrically. Taking into account that the whole-body metabolic rate scales more steeply in mammals than in birds, the mass-specific metabolic rate of visceral organs must decrease with body mass at a much faster rate in birds than in mammals. To explain this striking difference, there is an urgent need to study the metabolic rates of tissues and organs to supplement measurements of the whole-body metabolic rate.

## 1. Introduction

The slower than linear increase in metabolic rate with body mass, often referred to as a negative or sublinear or hypoallometric mass-scaling of metabolism, has fascinated biologists since at least the time of the publications of Rubner (1908) and Kleiber (1947). Technically, this phenomenon is studied by examination of the slope (*b*) in a linear regression, where log metabolic rate = *a* + *b* log body mass, which takes a value <1 under a hypoallometric scaling. Although the existence of a universal value for *b* has been contradicted in recent decades (e.g., Harrison 2018 and citations there), the ubiquity of the hypoallometric scaling requires explanation on two levels: ultimate and proximate. There is no agreement on the ultimate causes, whereas the proximate mechanism is clear but not always invoked: the size of metabolically relatively inert parts must increase with body mass faster than linearly, the size of energy-demanding organs must increase with body mass slower than linearly, the mass-specific metabolic rate of organs must decrease with body mass or, most likely, all of these phenomena occur simultaneously (Krebs 1950; Wang *et al.* 2001). Therefore, evidence of the interspecific scaling of organ masses is crucial for explaining the negative scaling of the whole-body metabolic rate on the proximate level.

Old data on scaling of organ masses in mammals has been summarized by Prothero (2015), who also performed his own analysis of original data extracted from the literature. However, his analysis, as well as analyses in older sources, did not take into account phylogenetic information, which is standard in contemporary research. Similarly, published analyses of body composition in birds (Peters 1983; Daan, Masman & Groenewold 1990) were phylogenetically non-informed.

Here, we used published data to estimate body mass-scaling for the mass of the brain, heart, liver, kidneys, lungs, spleen, digestive tract and its components (stomach and intestine) and the adipose deposits in 100 mammalian species and the mass of the brain, heart, liver, kidneys, lungs, breast muscle, skin, digestive tract, plumage and fat in 22 bird species with a phylogenetically informed method. We address two questions: the role of phylogenetic signal in organ/tissue size scaling and the potential role of organ/tissue mass scaling in interspecific metabolic rate scaling. Recent studies showed that hypoallometry of the metabolic rate is not an artefact of phylogeny, but taking into account phylogenetic information affects the slopes of the scaling (e.g., Griebeler & Werner 2016). Because the relative masses of organs may affect the scaling of the metabolic rate, it is important to examine the sensitivity of the slopes of organ masses to phylogenetic signal.

## 2. Methods

Data sets used in our analyses are relatively uniform, collected by the same people. Data for whole-body mass, fat-free body mass, adipose deposits and sizes of organs were taken from the supplemental material of Navarrete, van Schaik and Isler (2011). Their dataset comprises one species of Didelphimorphia, 3 species of Diprodontia, 3 species of Artiodactyla, 28 species of Carnivora, 3 species of Chiroptera, one species of Erinaceomorpha, two species of Lagomorpha, 23 species of Primata, 29 species of Rodentia, one species of Scandentia and 6 species of Soricomorpha. Data for 22 bird species were taken from Daan, Masman and Groenewold (1990). Their dataset comprises 9 species of Passeriformes, one species of Galliformes, 6 species of Charadriformes, one species of Columbiformes, one species of Falconiformes, 3 species of Anserifomes and one species of Rallidae. The species of birds were chosen by Daan, Masman and Groenewold (1990) to cover relatively uniformly log body mass axis. The authors also provide the BMR (basal metabolic rate) of the same birds that were used for tissue/organ mass analysis. Those BMR measurements were analysed here with the PGLS method here. Wet masses of mammalian and dry masses of birds’ organs were analysed; the slopes are comparable because water mass in birds scaled isometrically, whereas the intercepts are not. Phylogenetic trees of the studied birds and mammals are presented in the supplemental material (Figures S1 and S2).

Scaling parameters for sizes of organs/tissues were calculated in R software (R CoreTeam 2019) with an ordinary least squares regression (OLS) and with a phylogenetic generalized linear model (PGLS) from the caper package (Orme *et al.* 2018). Body mass or fat-free body mass were independent variables. All analysed data were log transformed prior to the analysis.

## 3. Results

The results of the PGLS and OLS models for mammals with log fat-free body mass as the independent variable are presented in Figure 1 together with the corresponding confidence intervals for the slopes (*b*). In mammals, the masses of the brain, heart, liver, kidneys, digestive tract as a whole, and intestine scale hypoallometrically for both the PGLS and OLS. Scaling of the spleen and stomach masses is hypoallometric according to the OLS, but PGLS analysis does not exclude isometry. Scaling of the lung and adipose deposit masses is isometric for both the OLS and PGLS. The results of the same analyses with log body mass are given in the supplemental material (Figure S3); because the increase in adipose deposits is almost ideally isometric (PGLS slope 0.99, OLS slope 1.02), the scalings for log body mass and for log fat-free body mass are almost identical. A substantial difference between the PGLS and OLS results is visible only in the scaling of the brain mass (PGLS slope 0.71, OLS slope 0.82), which demonstrates the strongest phylogenetic signal (Pagel’s lambda=0.923). Interestingly, the slightly weaker but still strong phylogenetic signals observed for the scaling of the stomach and spleen masses do not affect the slopes and only slightly affect the intercepts. The scaling of the liver, intestine and adipose deposit masses does not show any phylogenetic signals. The scattering of species-specific points around regression lines is very low (r^2^=0.97-98) in most visceral organs and low for brain (r^2^=0.94, PGLS), spleen (r^2^=0.81), stomach (r^2^=0.95) and adipose tissues (r^2^=0.88).

**Figure 1.**
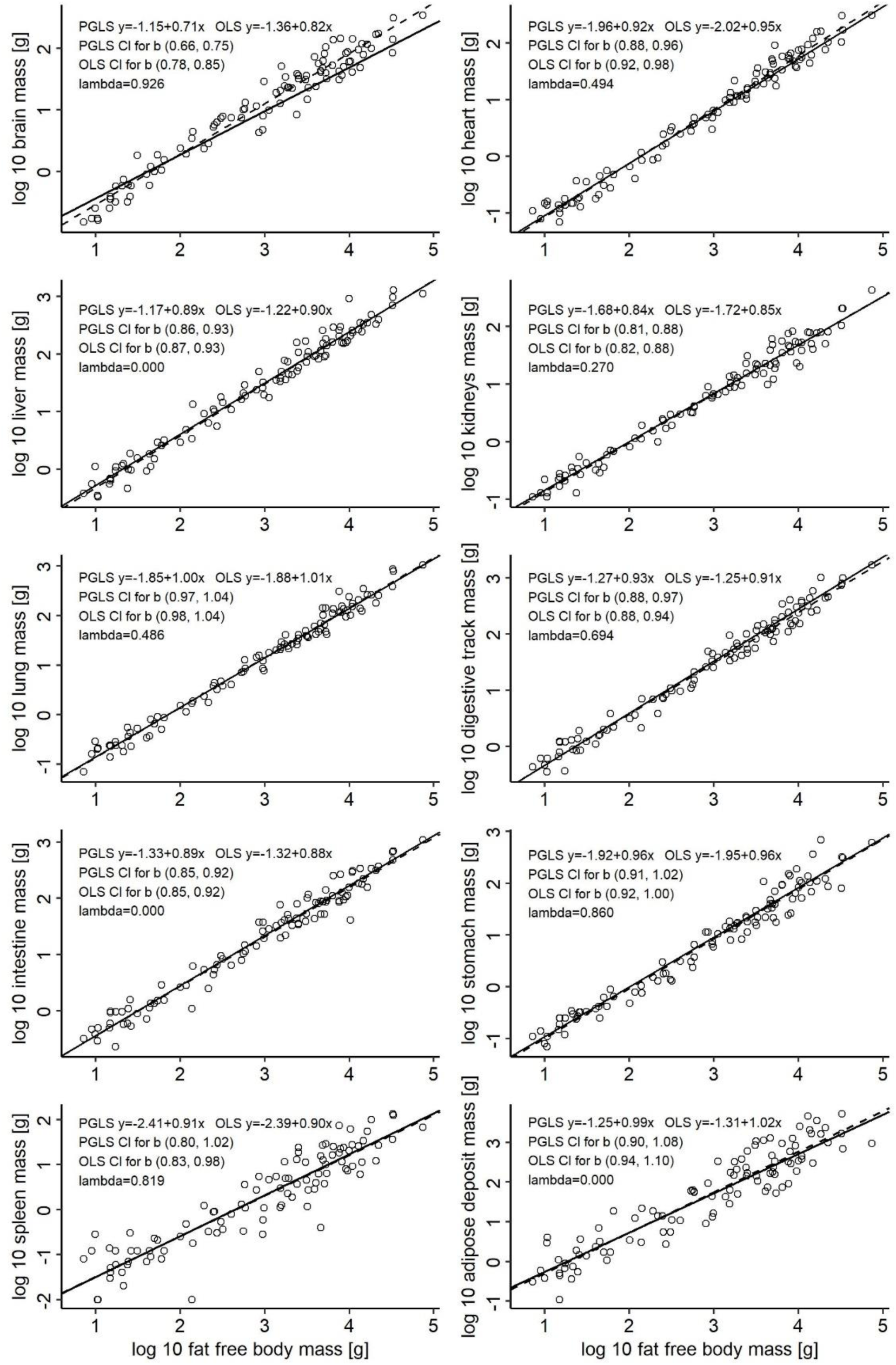
PGLS and OLS interspecific scaling of tissue/organ masses in mammals with log fat-free-body mass as the independent variable. For the scaling with log body mass as an independent variable, see the supplemental material, Figure S3.

In birds, water mass scales isometrically with body mass (the slope is 0.99 for both PGLS and OLS) with very narrow confidence intervals (Supplemental Figure S4). Thus, slopes for dry masses of organs in the studied birds are comparable to slopes for wet masses of organs in the studied mammals. Scattering of data points for fat mass in birds is substantial, with slopes of 0.95 for PGLS and 0.91 for OLS if fat-free body mass is taken as the independent variable (Figure 2). As a result, confidence intervals are very broad for this scaling, not excluding isometry. Thus, it is reasonable to show scaling with respect to fat-free body mass for other organs (Figure 2). Brain mass scales hypoallometrically with a slope of 0.69 (PGLS) or 0.56 (OLS) and a very strong phylogenetic signal. Breast muscle, heart, lung, kidney, liver and digestive tract masses scale isometrically or almost isometrically, with the isometric slopes falling within the confidence intervals. Phylogenetic signal is negligible for these organs. Plumage mass also scales isometrically but with a weak phylogenetic signal. Leg muscle mass scales isometrically but with a strong phylogenetic signal (Supplemental Figure S4). Interestingly, skin mass seems to scale hyperallometrically for OLS, but PGLS does not exclude isometry; phylogenetic signal is moderate for this tissue. Despite the much smaller number of species than in the case of mammals, r^2^ is very high, in the range of 0.93 to 0.99, with two exceptions: for fat mass, r^2^ equals 0.63 (PLGS) or 0.62 (OLS); for brain mass, r^2^ equals 0.93 in PGLS analysis but only 0.79 in OLS analysis, which additionally supports the need for phylogeny-informed analysis in the case of this organ.

**Figure 2.**
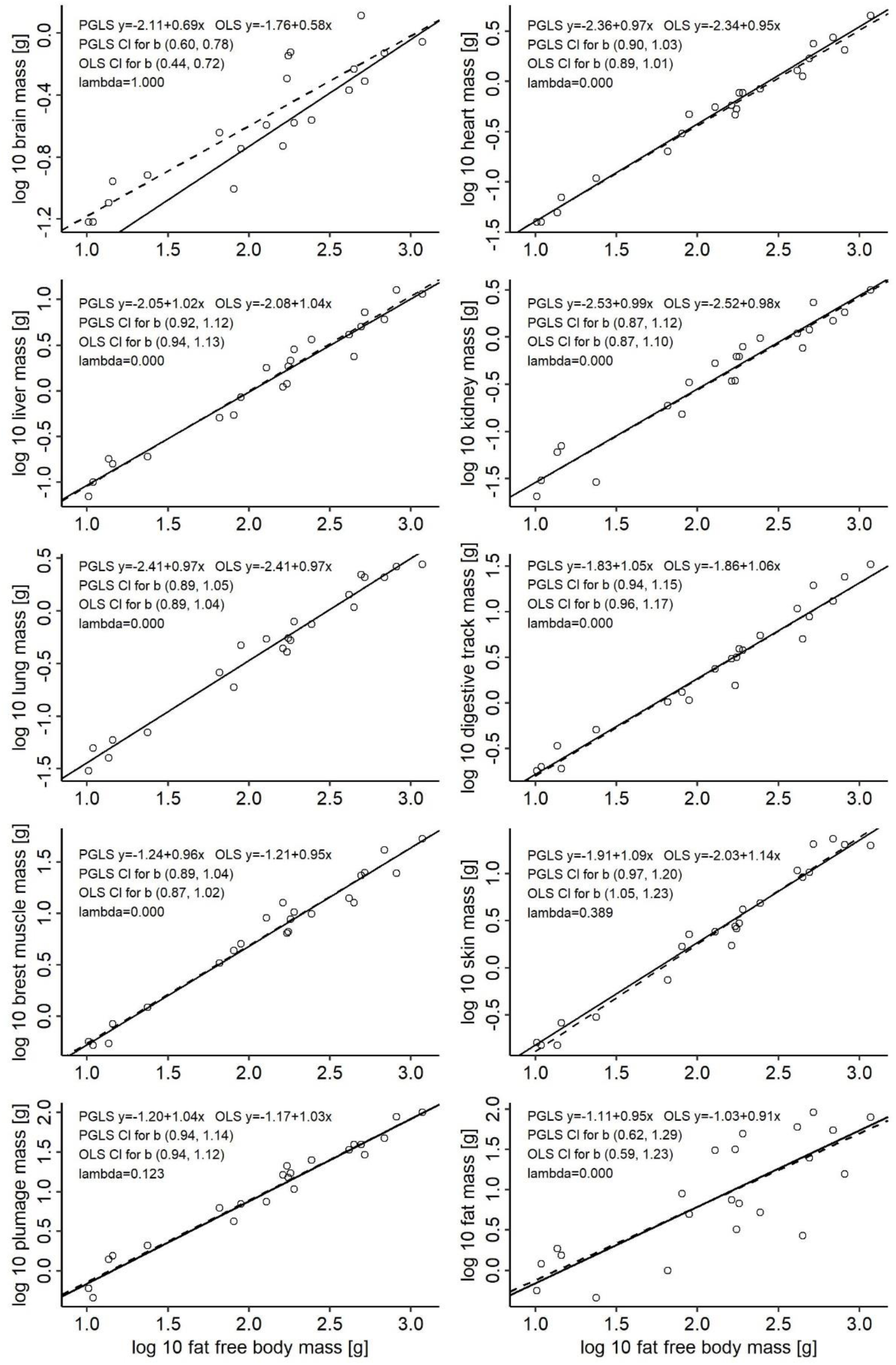
PGLS and OLS interspecific scaling of tissue/organ masses in birds with log fat-free-body mass as the independent variable. For the scaling with log body mass as an independent variable, see the supplemental material, Figure S5.

The slope for BMR in the studied birds equalled 0.67 according to PGLS analysis (Supplemental Figure S4), almost identical to the value reported by McKechnie and Wolf (2004). Thus, the set of birds analysed in this paper, although relatively small, represents very well the class of birds with respect to their metabolic requirements.

## 4. Discussion

Adipose tissue is metabolically relatively inert (1.90 kJ kg^-1^ h^-1^ in humans (Gallagher *et al.* 1998)), but our results show that its mass-scaling cannot explain the origin of the negative scaling of whole-body metabolism. We found that the mass of the adipose deposits scales with the log body mass isometrically in mammals (*b*=0.99 (PGLS) or 1.02 (OLS)) and almost isometrically in birds (*b*=0.95 (PGLS) or 0.91 (OLS)).

The lung mass scales isometrically with body mass in both mammals and birds. Prothero (2015) also reported isometric scaling of lung mass in 115 mammalian species from 12 orders. However, alveolar surface area scales with a slope of 0.94, pulmonary capillary surface area with a slope of 0.89 and diffusion capacity with a slope of 0.965 in 20 mammal species from 6 orders (Prothero 2015). According to Hoppeler and Weibel (1998), lungs in mammals have very low developmental and training plasticity, and they are built with significant excess structural capacity in non-athletic mammals but may limit the maximal metabolic rate in athletic species. Because of the high energy demand for flight, all studied birds can be considered athletic. Although Lasiewski and Calder (1971, cited after Peters 1983) reported a lower slope of the lung mass, equal to 0.95, in birds, it is likely that the confidence interval of this scaling contains isometry because r^2^ equals only 0.85 for their analysis. Additionally, the slope of 0.94 with SE 0.029 reported by Brody (1945) seems relatively close to isometry.

Our results show that the mass of the heart, responsible for the distribution of both oxygen and nutrients, has in mammals a slope of 0.92 (PGLS) or 0.95 (OLS). Prothero (2015) reported a slope 0.95 for 126 mammalian species from 14 orders. The slopes for birds in our analysis are slightly higher in the PGLS analysis (0.97), but confidence intervals contain mammalian values. Lasiewski and Calder (1971, cited after Peters 1983) reported a slope of 0.91 with r^2^=0.94 and Brody (1945) a slope of 0.92 with SE 0.021 in birds.

The liver mass scales in mammals with a slope of 0.89 (PGLS) or 0.90 (OLS) in our analysis, identical to the value given by Prothero (2015) for 134 species from 13 orders. Interestingly, the mass of liver scales isometrically in birds in our data, but Brody (1945) reported a much lower value 0.88 with SE 0.026. The slope for the kidney mass in mammals in our analysis, equal to 0.84 (PGLS) or 0.85 (OLS), is slightly lower than the 0.88 reported by Prothero (2015), but the difference fits the confidence interval. Again, in contrast to mammals, scaling of the kidney mass is isometric in birds. Previous papers reported lower slopes for the kidney in birds: 0.91 in 334 species (Johnson 1968) or 0.85 with the SE 0.032 (Brody 1945), likely because of including non-flying giants such as ostrich. The mass of the digestive tract as a whole scales hypoallometrically in mammals (0.93 for PGLS and 0.91 for OLS), while in birds, the scaling is even slightly hyperallometric: 1.05 (PGLS) or 1.06 (OLS), although confidence intervals contain isometry. Old data by Brody (1945) and Calder (1974, cited after Peters 1983) confirm isometric scaling.

Because of the expensive brain hypothesis, linking relative brain size with the life history-based pace of life (Aiello & Wheeler 1995), more is known about the scaling of the brain mass. The observed 0.70 PGLS slope in our analysis for mammals is lower and the 0.81 OLS slope is higher than the 0.77 slope (OLS, r=0.98; the same slope for 14 order averages) given by Martin (1996) for a set of 477 mammals. According to a recent analysis of 1552 species of mammals, the OLS slope for brain mass equalled 0.75 (CI 0.742, 0.758) with a very strong phylogenetic signal (Burger & George 2019); according to PGLS analysis, the slope was very low at only 0.57. However, the analysis on the level of orders resulted in variable slopes (0.24 to 0.81) with a median value of 0.64, differing from both OLS and PGLS slopes for all mammals. These results provide a warning that PGLS analysis does not solve the problem of the lack of phylogenetic independence if grade shifts exist, i.e., if branches of a tree differing in the slopes or intercepts of a scaling are not randomly distributed along the log body mass axis (Martin, Genoud & Hemelrijk 2005). Although the realistic value of the slope is still uncertain, the scaling of brain mass in mammals is much lower than one. It is also very low in birds in our study, as well as in Brody (1945), 0.498 with SE 0.022, and in Martin (1981), 0.58 SE 0.018. Because PGLS slopes for the brain are much higher than OLS slopes in both mammals and birds, old OLS slopes must be considered understated. However, even phylogeny-informed analysis gives slopes for the brain that are much lower than for other organs (0.71 in mammals and 0.69 in birds).

Plumage mass scales isometrically in the studied birds. Brody (1945) also reported isometric scaling of plumage in male passerine birds (slope 0.99, SE 0.130) and close to isometric in females (0.93, SE 0.098). Isometry of plumage mass is counterintuitive because larger birds have a smaller surface-to-volume ratio and thus lower heat loss. However, plumage also comprises flight feathers, and large birds are more likely to use gliding flight, requiring a greater bearing area. This explanation may also justify the hyperallometric scaling of skin mass found in our analysis, as skin is a base for anchoring flight feathers.

The heart, kidney, liver and brain are expensive organs. In humans, these organs comprise 0.5, 0.4, 2.6 and 2.0 % of the body mass, respectively, but are responsible for as much as 8.7, 8.2, 21.6 and 20.2 % of the total resting metabolic rate (calculated from Gallagher *et al.* 1998). Altogether, these organs constitute only 5.5 % of the body mass, but their metabolic rates constitute as much as 59 % of the metabolic rate (calculated from Gallagher *et al.* 1998). With isometric scaling (Raichlen *et al.* 2010; Muchlinski, Snodgrass & Terranova 2012), skeletal muscles constitute 40 % of body mass and use only 22 % of energy at rest in humans (Gallagher *et al.* 1998), and adipose tissue constitutes 21 % of body mass, using 15 % of the energy. Thus, the negative scaling of visceral organs responsible for 59 % of the metabolic rate in humans and over 50 % of that in mice (Konarzewski & Diamond 1995) may explain a large part of the negative scaling of the whole-body metabolic rate in mammals. Additionally, the mass-specific metabolic rate of these organs decreases with body mass, with a slope of -0.12 for the heart, - 0.27 for the liver, -0.08 for the kidneys (Wang *et al.* 2001), and -0.14 for the brain (Karbowski 2007). Interestingly, in birds, scaling of all visceral organs except the brain is steeper than that in mammals and isometric or close to isometric, but scaling of the whole-body metabolic rate is shallower (Glazier 2008). Because the brain mass of birds (1.7 % of body mass for 1 kg bird; Daan, Masman and Groenewold (1990)) is much lower than that of mammals (5.5 % of body mass for 1 kg mammal; Burger and George (2019)), shallower scaling of brain mass in birds cannot compensate for the steeper scaling of other visceral organs.

In birds, most tissues/organs scale isometrically. Strong hypoallometric scaling of the brain and slightly hypoallometric scaling of a few organs is likely to be balanced by hyperallometric scaling of the skin, plumage and digestive tract mass (if they truly diverge from isometry). In mammals, such hyperallometric scaling was not found. Since compensation must appear because body mass, by definition, scales isometrically with itself, hyperallometry may exist in tissues/organs not studied here, but we did not find strong enough hyperallometry in the survey presented by Prothero (2015) to support this view. Alternatively, some massive tissues/organs/components (water, fat, skeleton) considered to scale isometrically scale may in fact slightly hyperallometrically, which is possible considering the confidence limits for the scaling parameters.

The hypoallometric scaling of the masses of energy-demanding visceral organs must significantly contribute to the negative scaling of the whole-body metabolic rate in mammals, but such scaling only slightly contributes to the same scaling in birds. Taking into account that the whole-body BMR increases with body mass faster in mammals (PGLS slope between 0.71 and 0.74; White, Blackburn and Seymour (2009)) than in birds (independent contrast slope 0.68; McKechnie and Wolf (2004); 0.67 PGLS slope in the sample analysed here), the mass-specific metabolic rate of visceral organs must decrease with body mass much faster in birds than in mammals. Certainly, physiological mechanisms leading to this difference in proximate mechanisms shaping the metabolic rate scaling are worthy of study.

Metabolism takes place in cells that form different tissues/organs. Unfortunately, researchers usually measure total body masses and whole-body metabolic rates. Such state-of-the-art results do not originate from a well-developed research strategy but from the ease of taking these measurements. If we want to resolve the still-enigmatic hypoallometric scaling of the whole-body metabolism, we should refocus on body composition and organ-specific metabolic rates.

## Acknowledgements

We thank Karen Isler for providing the phylogenetic tree in a numerical form and Marcin Czarnoleski and Marek Konarzewski for comments on the manuscript.

## Supplemental material

### Phylogenetic trees of studied birds and mammals

**Figure S1.**
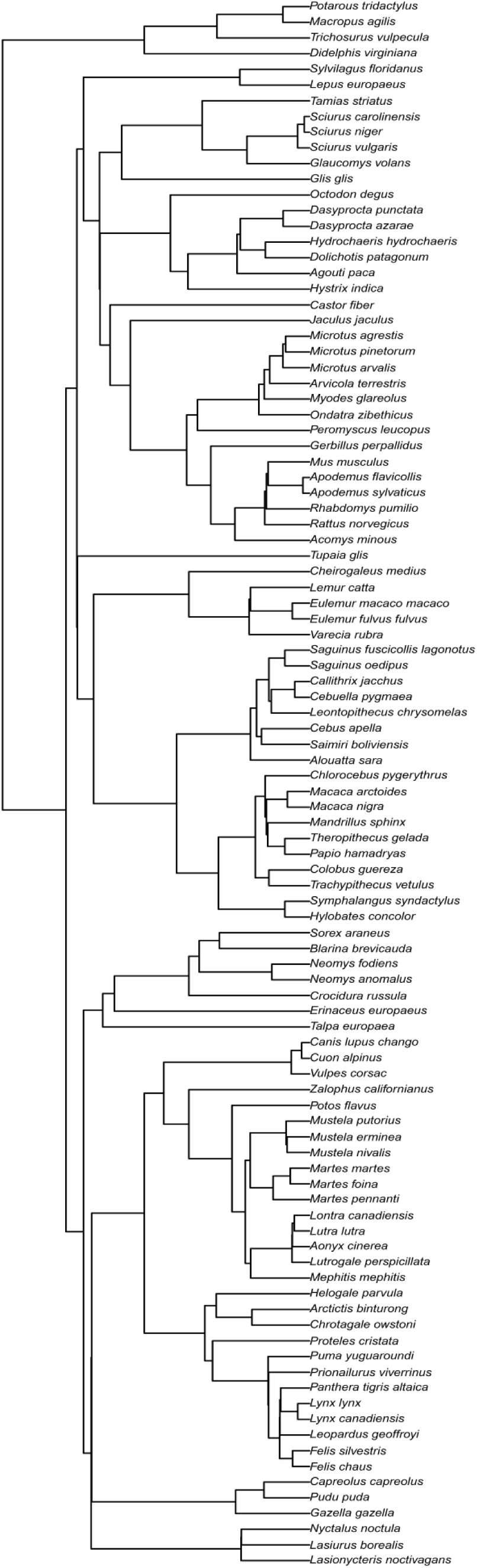
Phylogenetic tree of mammal species used in phylogenetically informed analysis. After: Navarrete, van Schaik and Isler (2011).

**Fig. S2.**
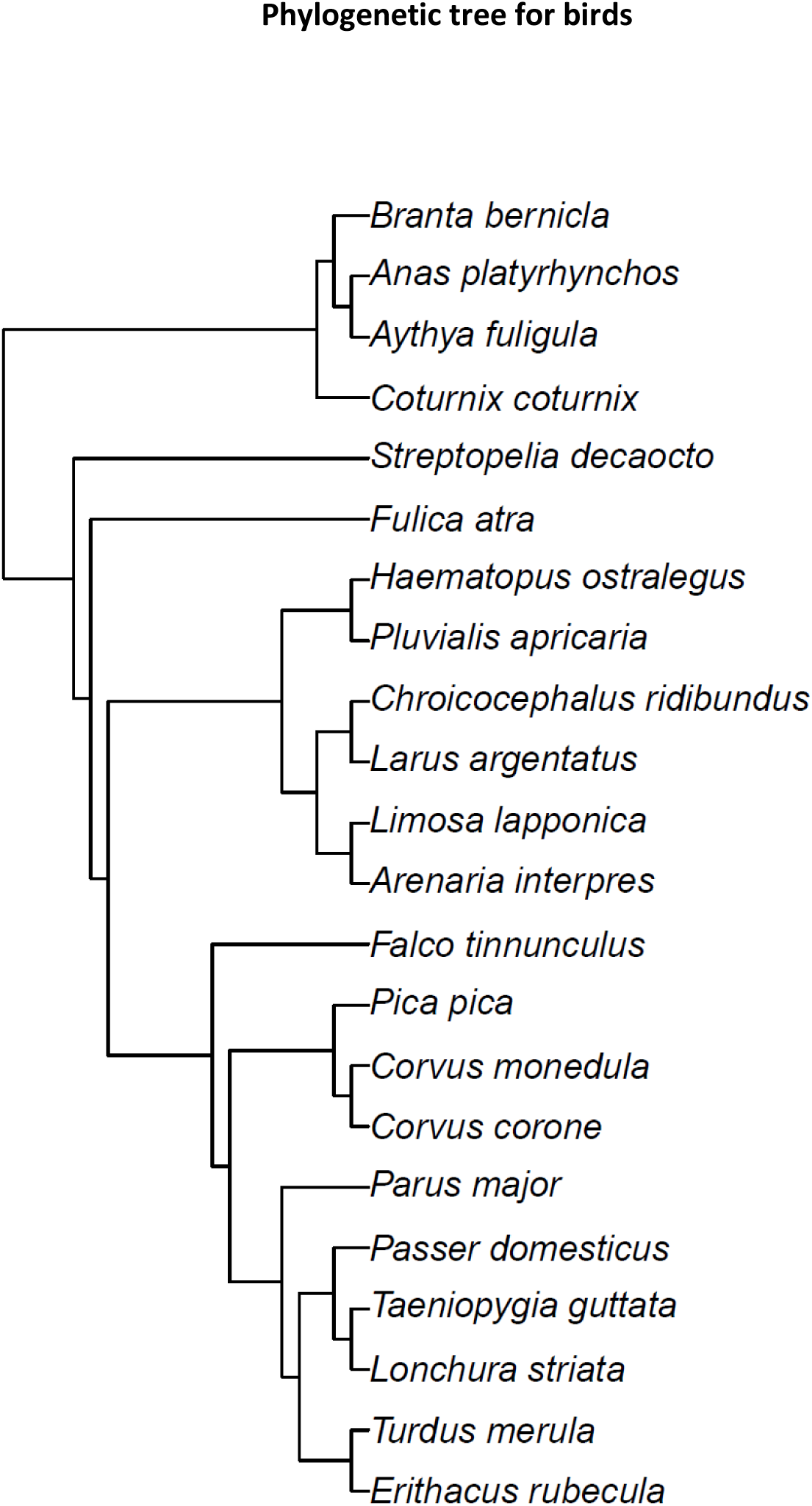
Phylogenetic tree of bird species used in phylogenetically informed analysis. Built with the data from Tree of Life: http://tolweb.org/tree/.

**Figure S3.**
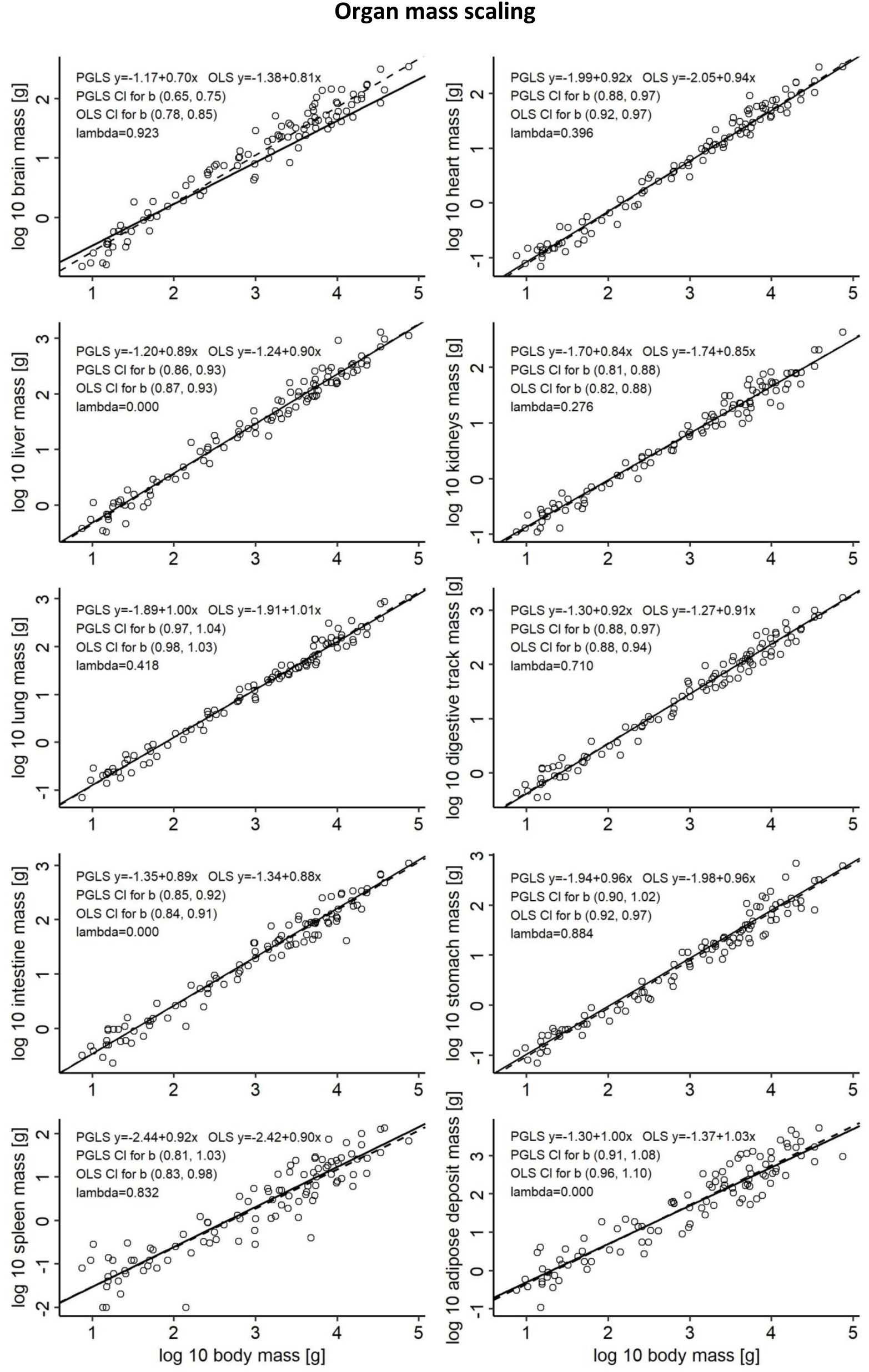
PGLS and OLS interspecific scaling of tissue/organ masses in mammals with log body mass as the independent variable.

**Figure S4.**
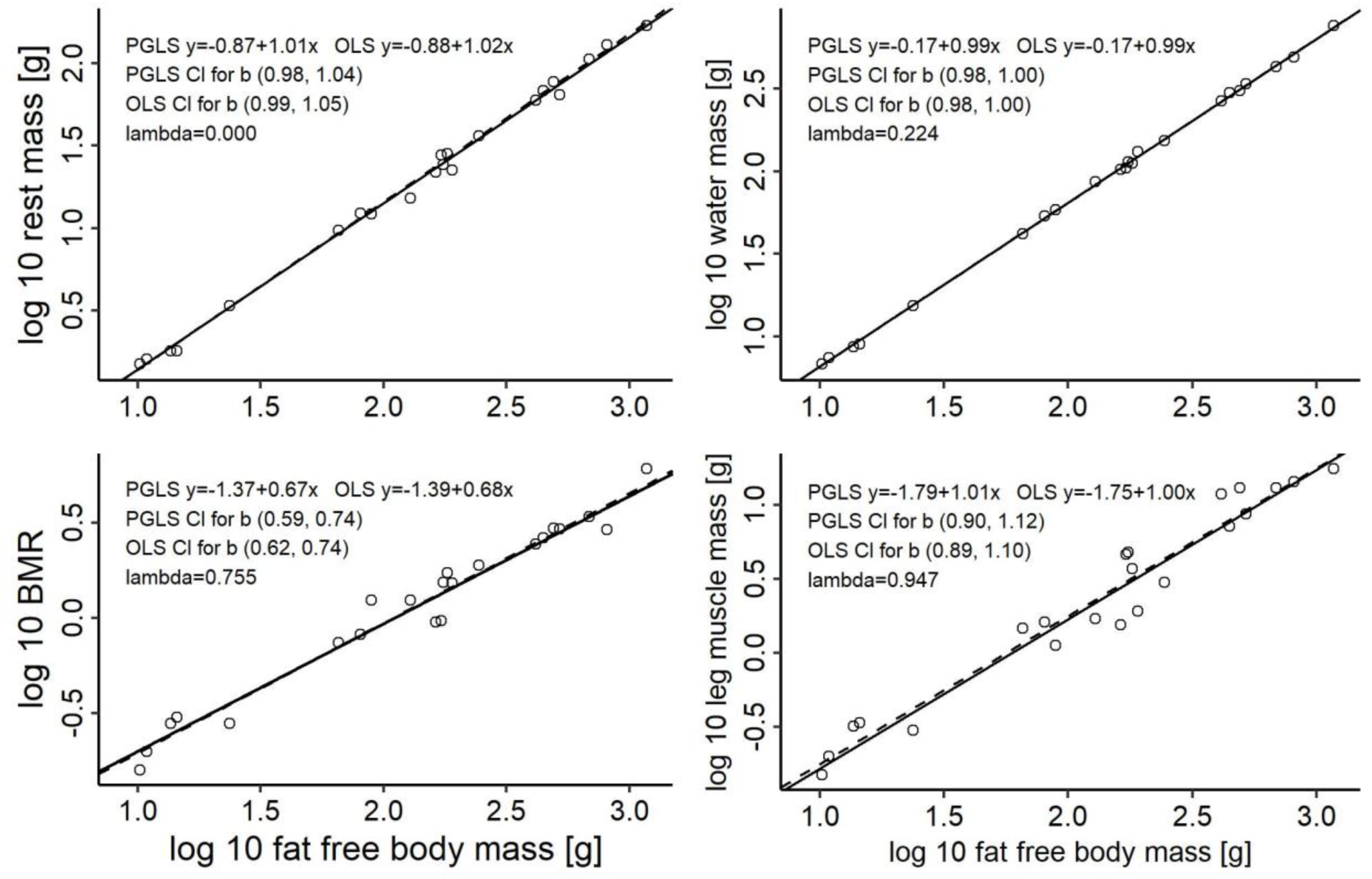
PGLS and OLS interspecific scaling of tissue/organ masses, water mass and BMR in birds with log10 fat-free body mass as the independent variable.

**Figure S5.**
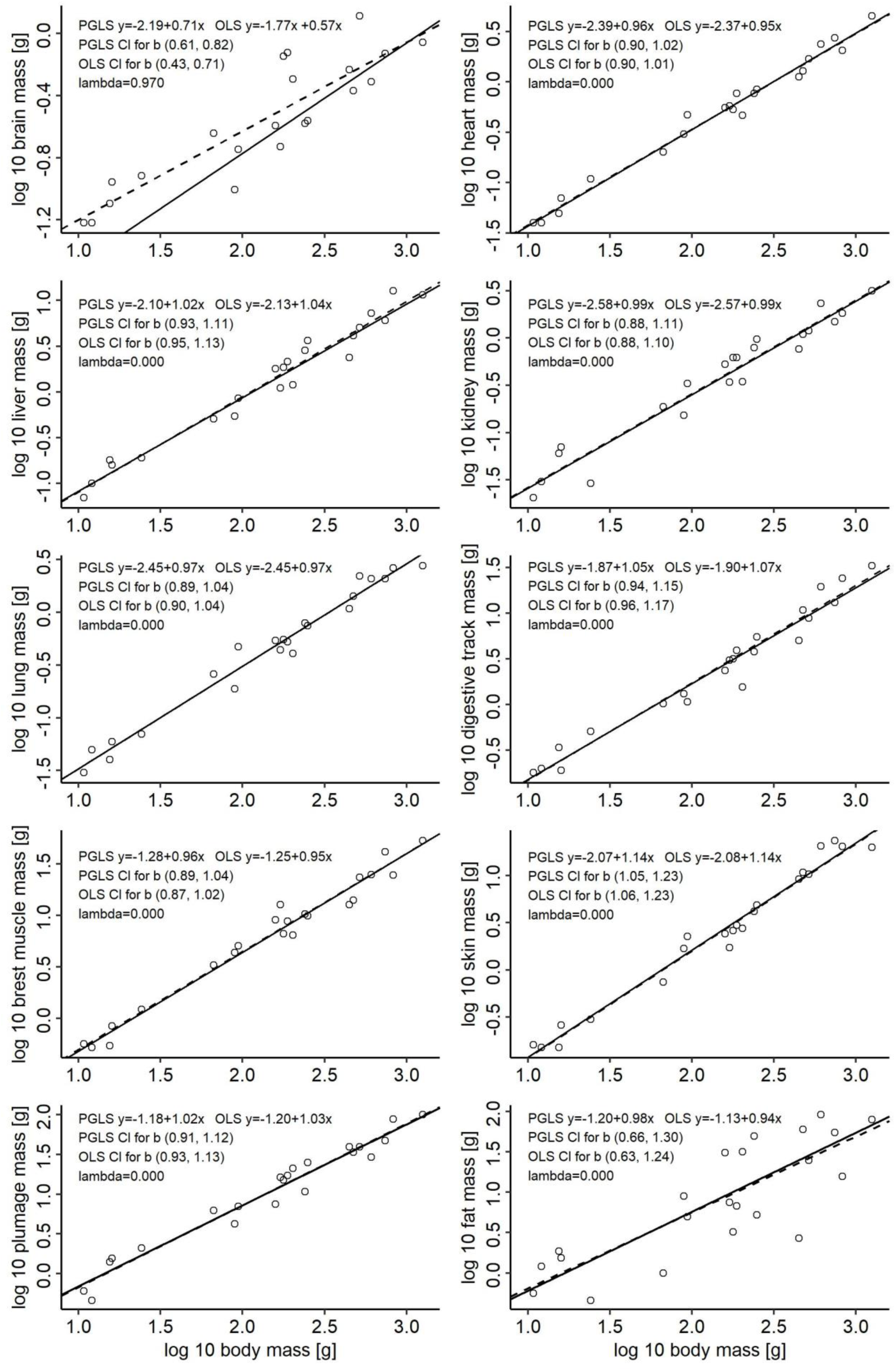
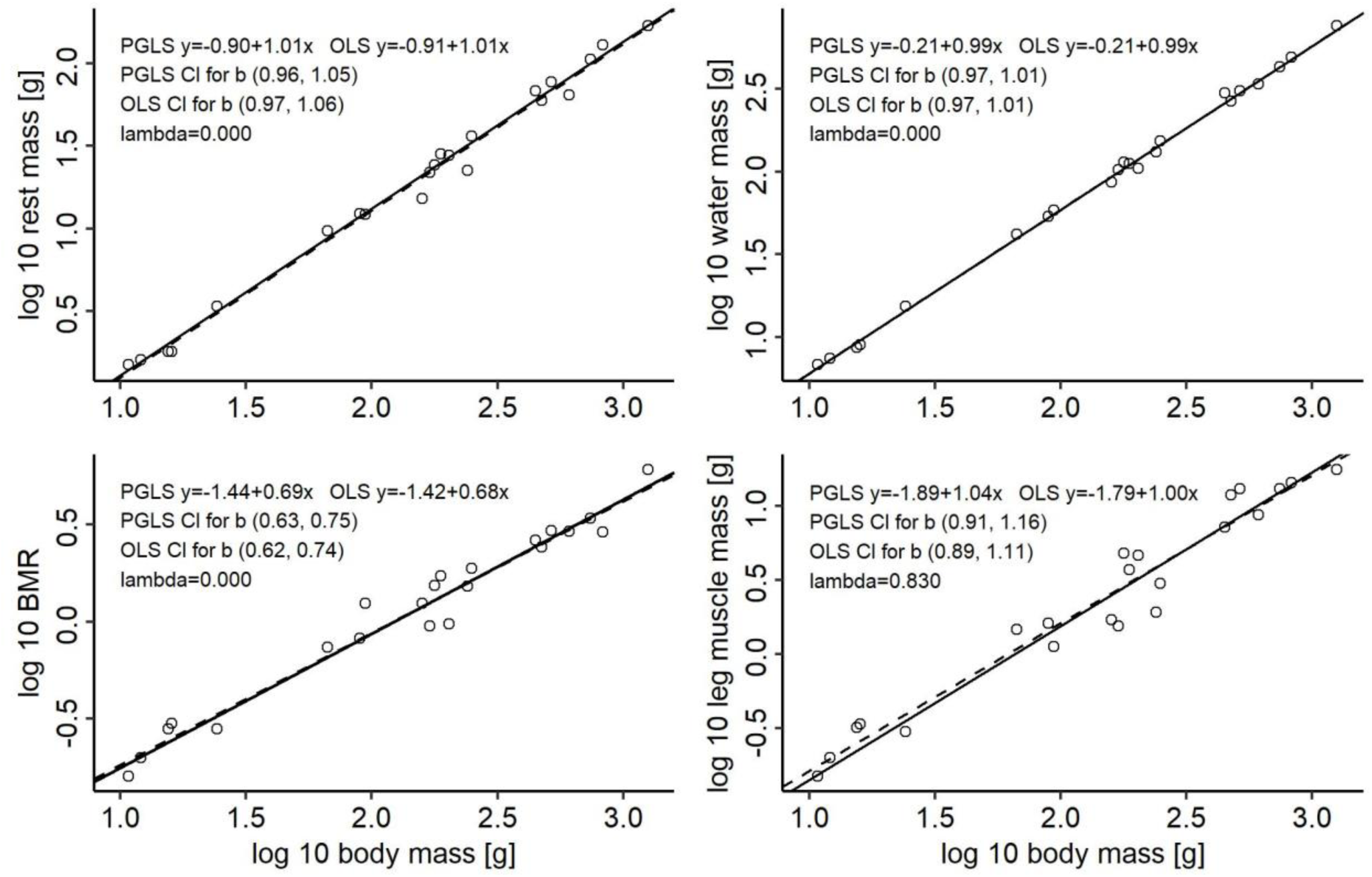
PGLS and OLS interspecific scaling of tissue/organ masses, water mass and BMR in birds with log body mass as the independent variable.

